# Interaction of contexts in context-dependent orientation estimation

**DOI:** 10.1101/816355

**Authors:** Ron Dekel, Dov Sagi

## Abstract

The processing of a visual stimulus is known to be influenced by the statistics in recent visual history and by the stimulus’ visual surround. Such contextual influences lead to perceptually salient phenomena, such as the tilt aftereffect and the tilt illusion. Despite much research on the influence of an isolated context, it is not clear how multiple, possibly competing sources of contextual influence interact. Here, using psychophysical methods, we compared the combined influence of multiple contexts to the sum of the isolated context influences. The results showed large deviations from linear additivity for adjacent or overlapping contexts, and remarkably, clear additivity when the contexts were sufficiently separated. Specifically, for adjacent or overlapping contexts, the combined effect was often lower than the sum of the isolated component effects (sub-additivity), or was more influenced by one component than another (selection). For contexts that were separated in time (600 ms), the combined effect measured the exact sum of the isolated component effects (in degrees of bias). Overall, the results imply an initial compressive transformation during visual processing, followed by selection between the processed parts.

**Highlights:**

- Non-linear sub-additivity for increased context area or contrast
- Non-linear selection between overlapping or adjacent, dissimilar contexts
- Linear additivity for combinations of temporally separated contexts

## Introduction

Visual processing is continuously adjusted based on previous visual stimulations, a process termed visual adaptation (Clifford et al., 2007; Kohn, 2007; Webster, 2015). Similarly, the processing of a visual region can be influenced by the surrounding visual stimulation (Lamme, 1995; Sagi, 1995; “The tilt illusion: Phenomenology and functional implications,” 2014). Changes in processing due to context are potentially useful, allowing for adjusting the dynamic range, calibrating transmitted information, and enhancing spatial and temporal discontinuities. Correspondingly, context, both temporal and spatial, gives rise to salient perceptual effects, often referred to as visual illusions (Clifford & Rhodes, 2005; Gibson, 1937; Gibson & Radner, 1937; Webster, 2011).

Context may affect both sensitivity and appearance. In the case of orientation features, the influence of a surrounding region (“spatial context”) on appearance can be measured by the tilt illusion (TI, Clifford, 2014; Gibson, 1937; Fig. 1A), by effects on contrast sensitivity (Cannon & Fullenkamp, 1993; Polat & Sagi, 1994a, 1994b, 1993), and by effects on orientation sensitivity (JND; Solomon & Morgan, 2006). The influence of a previous time frame (“temporal context”) can be measured by the tilt aftereffect (TAE, Gibson & Radner, 1937; Fig. 1B), by effects on contrast sensitivity (Blakemore & Campbell, 1969; Tanaka & Sagi, 1998), and by effects on orientation sensitivity (JND; Regan & Beverley, 1985). There are clear qualitative similarities between space and time (Schwartz, Hsu, & Dayan, 2007). In both cases, the oriented context may lead to a change (bias) in appearance, such that the perceived orientation is shifted in the opposite direction from the context orientation (though for very large orientation differences between context and target, the direction of the effect is inverted, which is known as the “indirect effect”, Clifford, 2014; Schwartz et al., 2007). The context induced perceived orientation shifts are often, but not always, accompanied by reduced contrast (Regan & Beverley, 1985) or orientation (Solomon & Morgan, 2006) sensitivity, with the relationship between the different effects depending on spatial and temporal stimulus configuration (Webster, 2015), and on experience (Chen & Fang, 2011). The stimulus manipulations used in the present study affected the bias in perceived orientation (TAE/TI) but not so much, if at all, the sensitivity to orientation differences (JND).

**Fig. 1.**
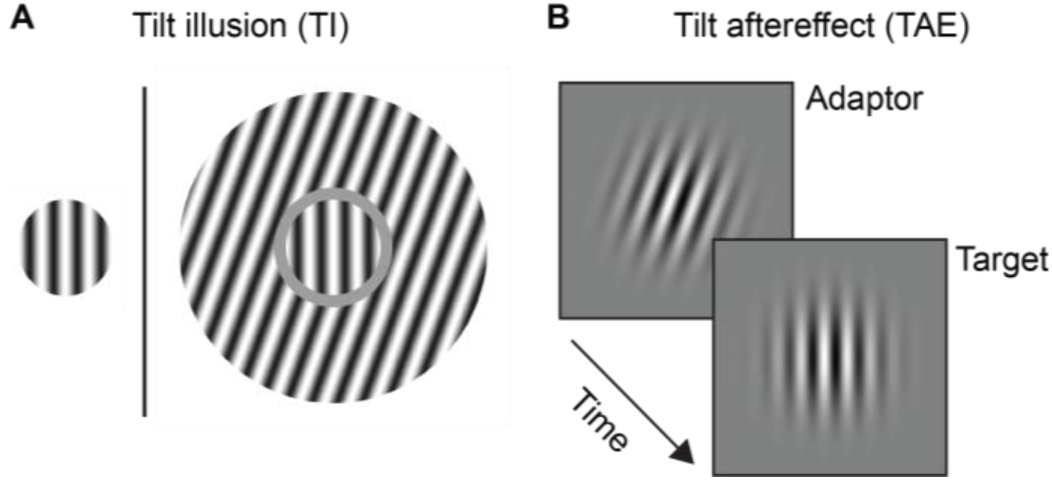
Tilt illusion and the tilt aftereffect. (**A**) In the tilt illusion (TI), an oriented surround (here, an annulus) leads to a change in the perceived orientation of a target (here, the center circle). The target without surround, shown on the left, is provided as a reference and was not used in the experiments. (**B**) In the tilt aftereffect (TAE), exposure to an oriented adaptor leads to a change in the perceived orientation of a subsequently viewed target. Figure reproduced from (Dekel & Sagi, 2020).

Traditionally, contextual effects on perceived orientation were thought to be mediated by shifts of internal “norms” for vertical and horizontal orientations (Gibson & Radner, 1937). More recent models consider this calibration process to be stochastic (Solomon & Morgan, 2006). According to other theories, bias may reflect early visual adaptation (in time) and lateral inhibition (in space) of orientation-selective units (Blakemore & Campbell, 1969; Blakemore, Carpenter, & Georgeson, 1971; Coltheart, 1971). However, more recent evidence implies the involvement of more complex mechanisms, such as learning of statistical regularities (Nakashima & Sugita, 2014; Pinchuk-Yacobi, Dekel, & Sagi, 2016), task experience (Dong, Gao, Lv, & Bao, 2016; Pinchuk-Yacobi, Harris, & Sagi, 2016; Yehezkel, Sagi, Sterkin, Belkin, & Polat, 2010), and perceptual grouping (He, Kersten, & Fang, 2012; Suzuki, Clifford, & Rhodes, 2005). Such observations are important because natural vision is clearly more complex than the testing conditions of most laboratory experiments, raising the question of how findings about context obtained with reduced stimuli can be extended to actual natural stimulation (Schwartz, Snow, & Coen-Cagli, 2017; Solomon & Kohn, 2014). Specifically, most research about the TAE and TI is performed using simplified stimuli, such as lines (Gibson, 1937; Gibson & Radner, 1937), gratings (Campbell & Maffei, 1971; Mitchell & Muir, 1976), or Gabor patches (Knapen, Rolfs, Wexler, & Cavanagh, 2010). However, natural stimulation has high-order statistical regularities in space and time, leading to differences in the operation of the visual system for simplified vs. natural input (Alam, Vilankar, Field, & Chandler, 2014; Carandini et al., 2005; Movshon & Simoncelli, 2014; Olshausen & Field, 2005). It is therefore very important to understand how contextual influence differs between ecological vision and the laboratory (Coen-Cagli, Kohn, & Schwartz, 2015; Schwartz et al., 2017; Solomon & Kohn, 2014).

Recently, we used unmodified natural images that were selected to match the oriented low-level properties of the reduced stimulation, and found TAE due to image exposure (Dekel & Sagi, 2015; see also, Goddard, Clifford, & Solomon, 2008; Ismail, Solomon, Hansard, & Mareschal, 2016). This finding is consistent with a simple view of contextual influence: that it follows the sum over all the orientation energy in the image, regardless of a higher-order structure. Here, to address this simplified hypothesis in a systematic fashion, we considered different ways of combining contexts (in space/time, overlapping/non-overlapping), and compared the bias from a combined context, to the sum of the biases from the isolated components (additivity). This is a crucial first step towards more complex combinations, and natural vision.

Specifically, we performed the following experiments (overviewed at Fig. 2):

**Fig. 2.**
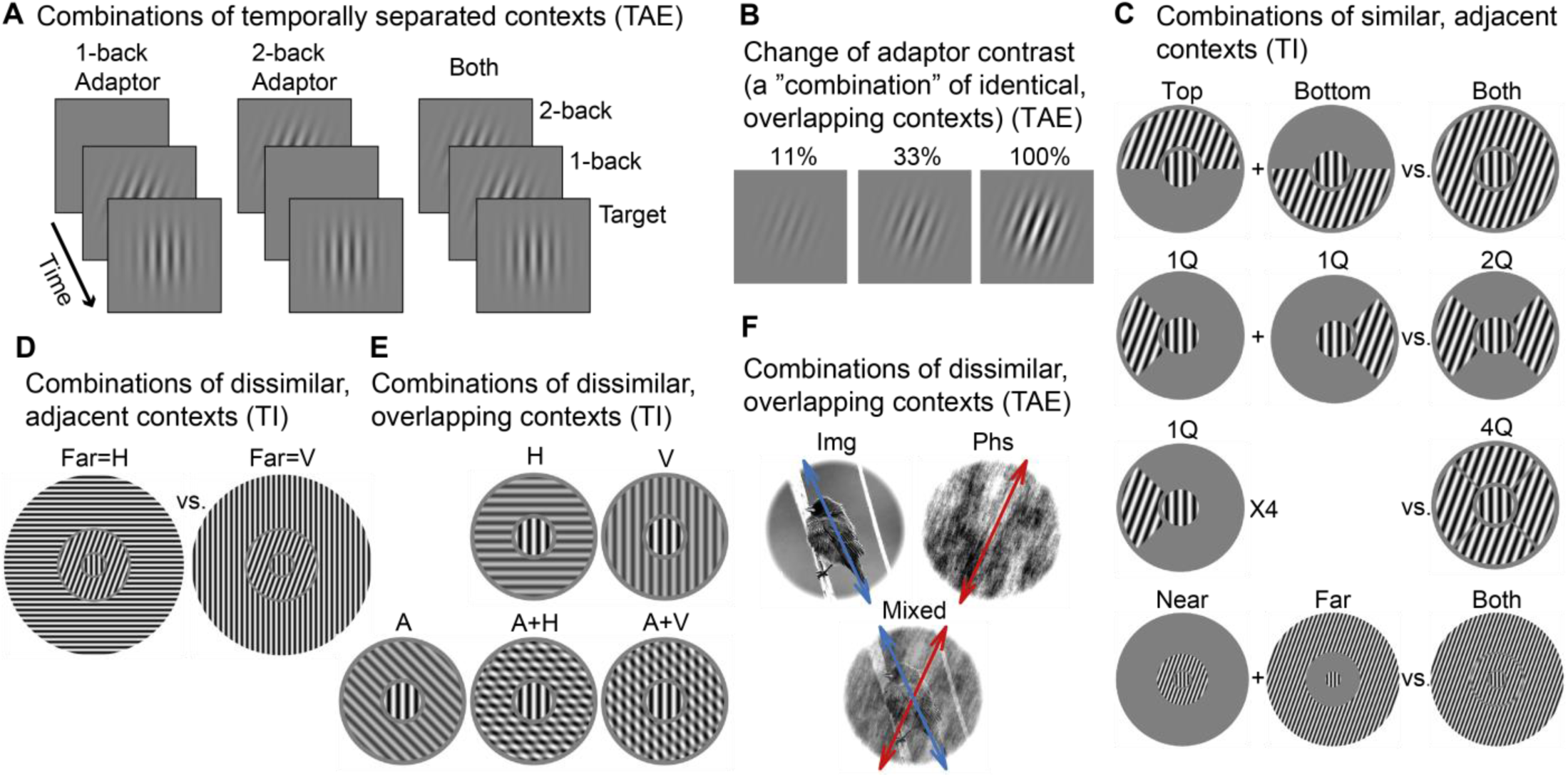
Summary of the main experimental designs in this work. Results presented at (**A**) Fig. 4, (**B**) Fig. 5, (**C**) Fig. 6, (**D**) Fig. 7, (**E**) Fig. 8, and (**F**) Fig. 9.

- First, we investigated the additivity of TAE for combinations of contexts presented at different times (separated). The results showed additivity, under both fixation and periphery conditions. This is the main finding of this work. To account for the additivity, we offer an explanation in terms of an optimal combination of independent reference cues (see Figs. 3 and 4, as well as the Discussion).
- In addition, we considered experiments with different adaptor contrasts, which showed sub-linearity of the TAE vs. contrast function (Fig. 5). This finding rules out the possibility that the additivity of separated contexts is due to measuring a TAE with a small magnitude.
- Next, we investigated the additivity of combinations of similar contexts presented at the same time. The influence of such combinations can be measured at the same (TI) or at a later (TAE) time. Here we only report results for TI, because TAE magnitudes for non-retinotopic combinations were too small (data not shown). The results for TI generally showed sub-additivity (Fig. 6), and interestingly, suggest spatial anisotropy of the sub-additivity. To interpret findings of sub-additivity (Figs. 5 and 6), we considered compressive transformation of information during visual processing (see the Discussion).
- Finally, we investigated combinations of dissimilar contexts. Such experiments are important, because natural stimuli are composed of dissimilar orientations. We considered combinations that are adjacent in space (TI, Fig. 7), overlapping (i.e., plaids, using TI, Fig. 8), or that use natural images (TAE, Fig. 9). The results generally showed non-linearity, showing a strong dependence on the spatial arrangement of the components. To interpret these findings, we considered selection between competing contexts (see the Discussion).

**Table 1.** Procedural information.

|  | #Obs.<br>used | #Obs.<br>pruned | Sessions<br>per obs. | Blocks per<br>session | Trials per<br>block | Total<br>used<br>trials | Total<br>pruned<br>trials | Fig. |
| --- | --- | --- | --- | --- | --- | --- | --- | --- |
| 1- and 2-back fixation | 9 | 0 | 1-2 | 5 | 93 | 4,633 | 17 | 4BDE |
| 1- and 2-back periphery<br>(AAAAT) | 5 | 0 | 5-10 | 5 | 65 | 13,271 | 39 | 4CDF |
| 1- and 2-back periphery<br>(AT/AAT/AAAT/AAAAT) | 6 | 1 | 7-9 | 5 | ~95<sup>(**)</sup> | 15,608 | 15 | 4CDF |
| Contrast Gabor | 9 | 0 | 1 | 5 | 125 | 5,560 | 65 | 5 |
| Contrast ‘biased’ images | 10 | 0 | 1 | 4 | 102 | 4,077 | 3 | 5 |
| Area near/far 400ms | 21<sup>(*)</sup> | 7 | 1 | 4 | 228 | 19,119 | 33 | 6DG |
| Area top/bottom 100ms | 19<sup>(*)</sup> | 7 | 1 | 4 | 228 | 17,293 | 35 | 6EH |
| Area top/bottom 400ms | 26<sup>(*)</sup> | 3 | 1 | 4 | 228 | 23,670 | 42 | 6EH |
| Area quadrants Q1/Q2 100ms | 20<sup>(*)</sup> | 5 | 1 | 4 | 228 | 18,220 | 20 | 6FI |
| Area quadrants Q1/Q2 100ms,<br>onset jitter | 13<sup>(*)</sup> | 7 | 1 | 4 | 228 | 11,806 | 50 | 6FI |
| Area quadrants Q1/Q4 100ms | 10<sup>(*)</sup> | 5 | 1 | 4 | 228 | 9,118 | 2 | 6FI |
| Area quadrants Q1/Q4 400ms | 17<sup>(*)</sup> | 5 | 1 | 4 | 228 | 15,489 | 15 | 6FI |
| Far=V/H, 100ms | 9<sup>(*)</sup> | 6 | 1 | 4 | 190<sup>(***)</sup> | 5,444 | 28 | 7 |
| Far=V/H, 400ms | 14<sup>(*)</sup> | 14 | 1 | 4 | 190<sup>(***)</sup> | 8,510 | 2 | 7 |
| Far=H/V/E, 100 ms | 26<sup>(*)</sup> | 6 | 1 | 4 | 228 | 23,629 | 83 | 7 |
| Far=H/V/E, 400 ms, onset jitter | 15<sup>(*)</sup> | 1 | 1 | 4 | 228 | 13,655 | 25 | 7 |
| Near=45°, Far=V/H, 400ms | 12<sup>(*)</sup> | 11 | 1 | 4 | 190<sup>(***)</sup> | 7,279 | 17 | 7 |
| 2nd order 400ms | 10 | 0 | 1 | 4 | 228 | 9,119 | 1 | 8 |
| 2nd order 400ms | 7<sup>(*)</sup> | 2 | 1 | 4 | 228 | 6,384 | 0 | 8 |
| 2nd order 100ms | 12<sup>(*)</sup> | 2 | 1 | 4 | 228 | 10,896 | 48 | 8 |
| Img+Phs common | 9<sup>(*)</sup> | 3 | 1 | 4 | 102 | 3,635 | 37 | 9 |
| Img+Phs laboratory | 10 | 0 | 1 | 4 | 102 | 4,079 | 1 | 9 |
| Img+Phs color | 10<sup>(*)</sup> | 0 | 1 | 4 | 102 | 4,072 | 8 | 9 |
| Img+Phs 100ms | 11<sup>(*)</sup> | 2 | 1 | 4 | 102 | 4,484 | 4 | 9 |
| Img+Phs detection task | 19<sup>(*)</sup> | 6 | 1 | 4 | 102 | 7,749 | 3 | 9 |

315 (\*) *MTurk observers.*
(\*\*) *In the “AT/AAT/AAAT/AAAAT” experiment, ~25% of the trials were “AT”, i.e., adaptor-test trials lacking a 2-back adaptor, and were ignored in the analysis. The remaining trials had either two, three, or four adaptor presentations preceding the test, denoted by AAT, AAAT, and AAAAT, respectively, with ~25% of the trials belonging to each sequence type.*
320 (\*\*\*) *In the “Far=V/H” experiments, 20% of the trials had an orientation of 0° or 90° (V or H) in both near and far surround annuli, and were ignored in the analysis.*

**Fig. 3.**
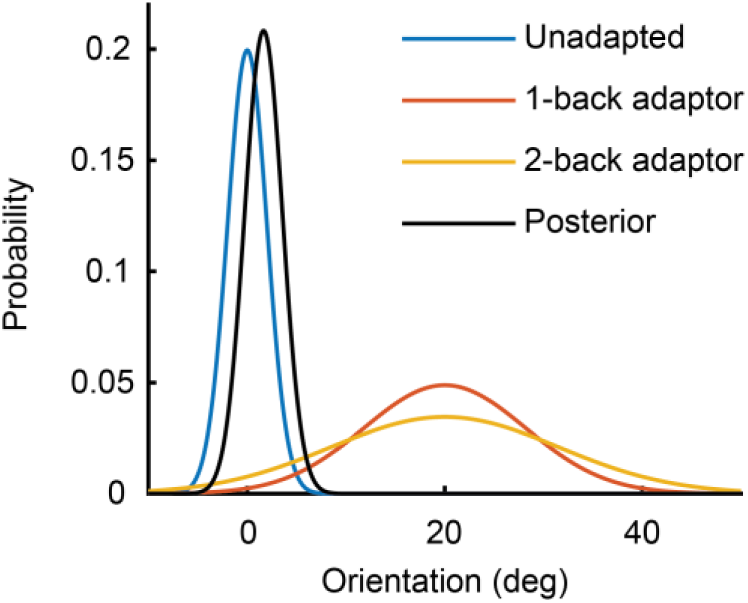
Illustration of the cue combination. The reference frame (i.e., posterior) is shifted in the direction of the adaptor cues, leading to a shift of the perceived orientation in the opposite direction.

**Fig. 4.**
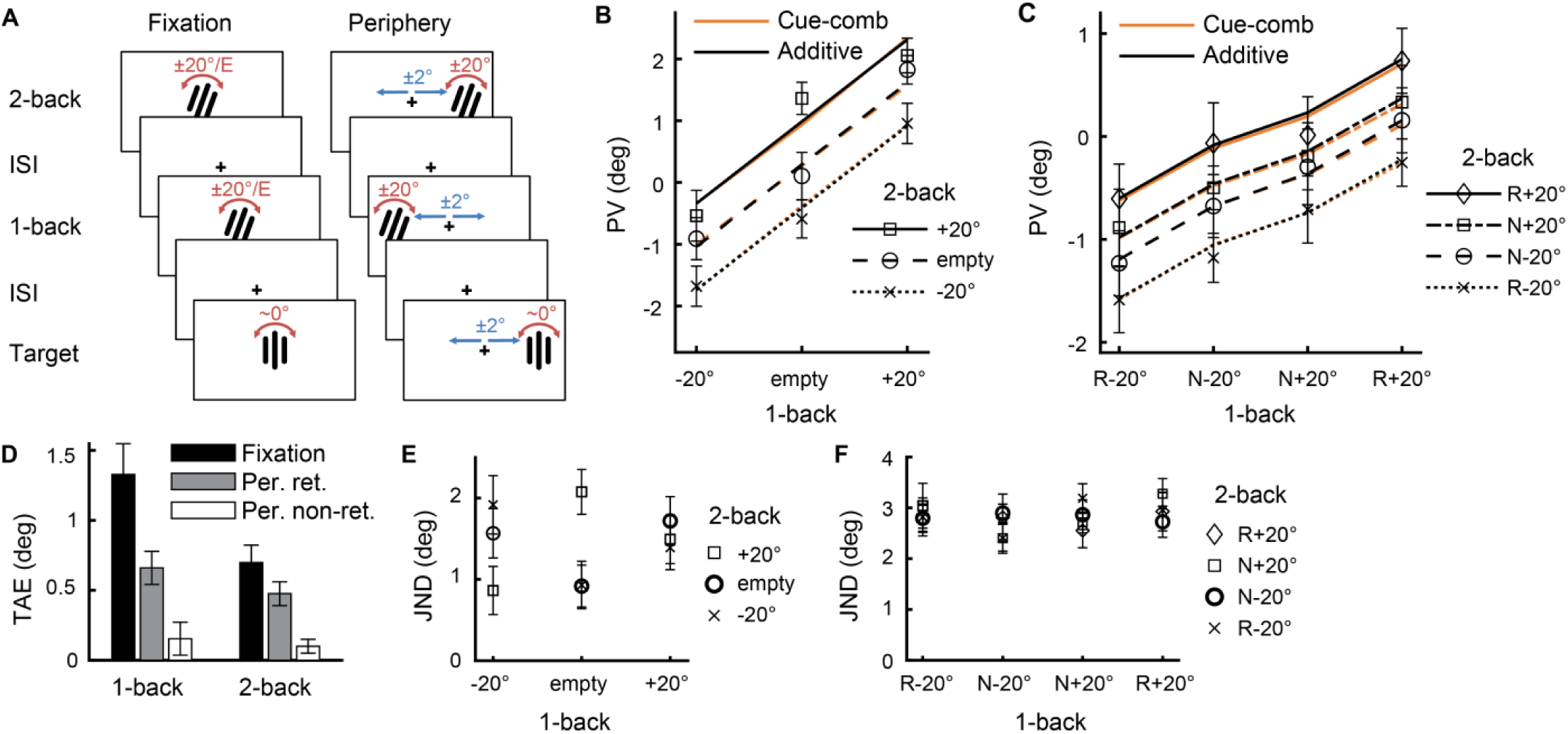
Additivity in temporal combinations of contexts. (**A**) Stimuli illustrations (actual stimuli were Gabor patches, see Fig. 1B). (**B and C**) Data points are the average across observers of the perceived vertical orientation (PV), measured for the “Target”, following exposure to different combinations of 1- and 2-back adaptors. The shown lines are the predictions based on additivity (black, Eqs. (1) and (2), using the averages shown in panel D) or the cue combination model (orange, Eq. (3)). (**B**) Fixation experiment. (**C**) Periphery experiment, where each of the 1- and 2-back adaptors was either presented at the same side as the target (retinotopic, “R”) or at the contra-lateral side (non-retinotopic, “N”). Note that the 1-back, 2-back, and Target locations were each randomized independently (Left/Right). The results showed linear additivity across all conditions. (**D**) Average TAE magnitudes. Note that magnitudes of TAE and TI, as shown in the subsequent figures, represent the change in PV. (**E and F**) Just noticeable differences (JND, see Methods) of the orientation discrimination task, for the (**E**) Fixation and (**F**) Periphery experiments. Error bars are ±1SEM.

**Fig. 5.**
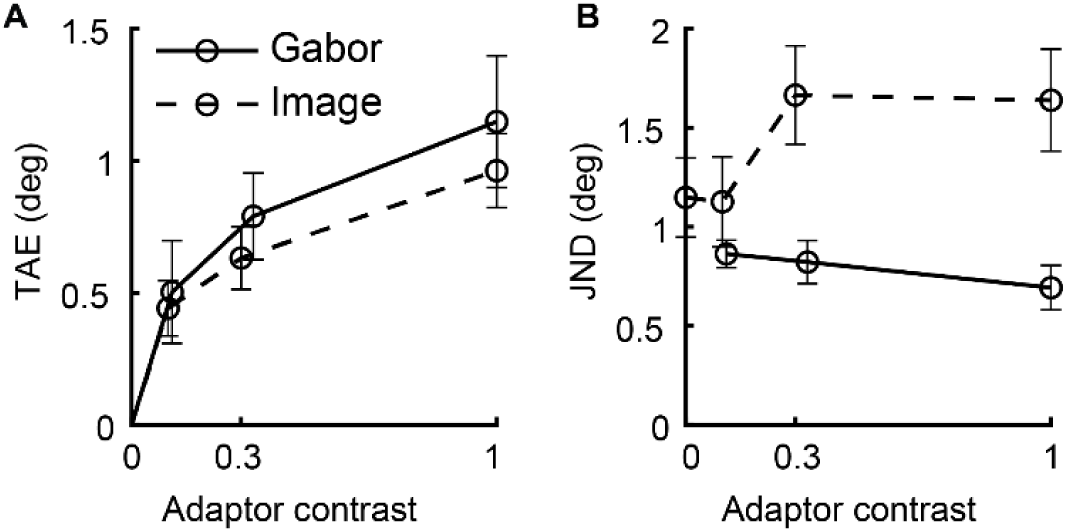
Sub-linearity of bias for increased adaptor contrast. (**A**) The average TAE across observers measured when using different adaptor contrasts is shown. TAE was caused by exposure to either a “Gabor” patch or an oriented “Image” adaptor, and measured using a near-vertical Gabor target (the “Gabor” case is comparable to Fig. 4). For contrast=0, we have TAE=0 by definition. The results show the sub-linearity of the TAE vs. contrast function. (**B**) Same, for JND. Error bars are ±1SEM.

**Fig. 7.**
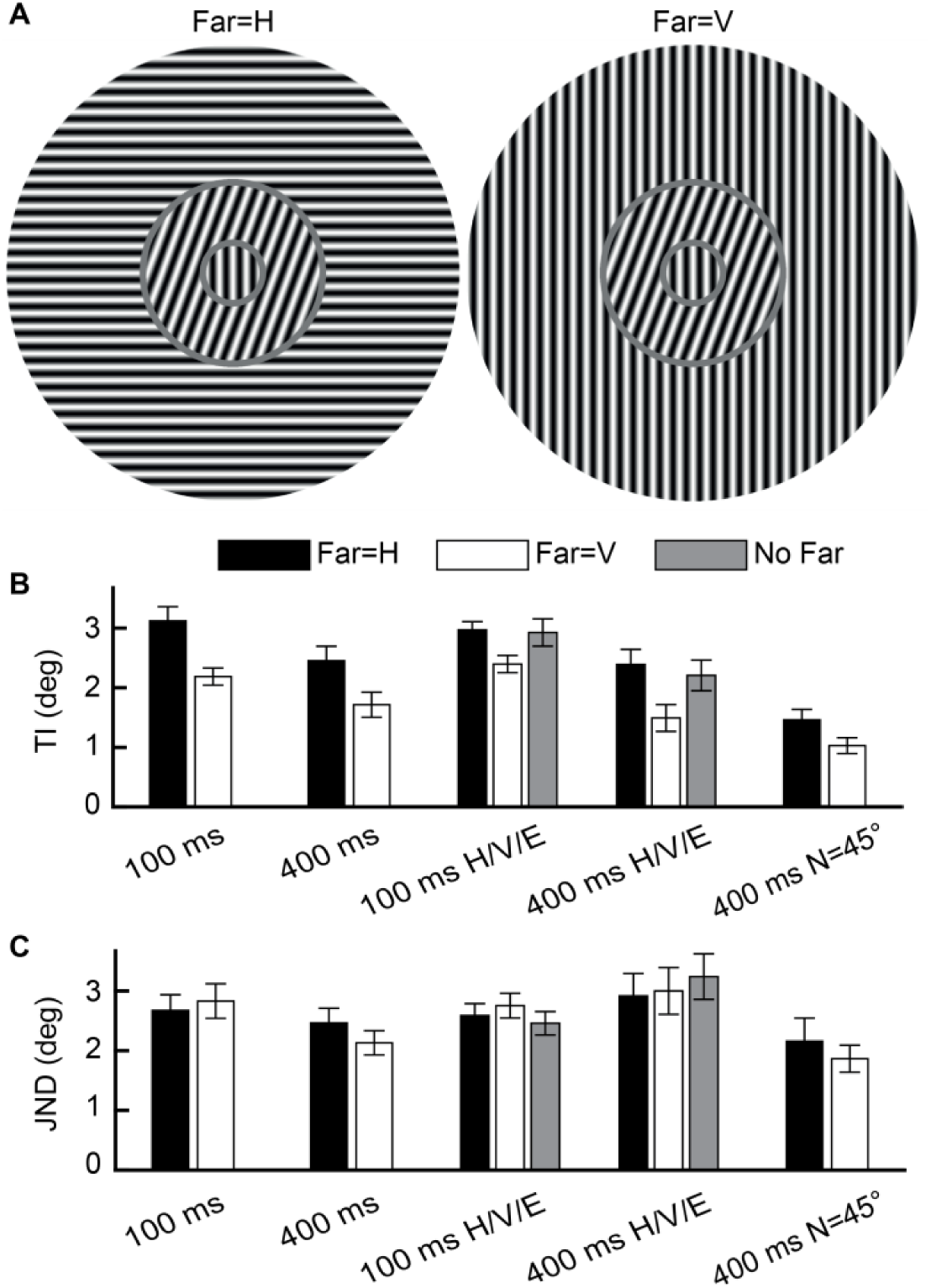
Non-linear competition between lateral interactions. (**A**) Stimuli illustrations. (**B**) Average across observers of the TI due to a near-surround annulus, when a far-surround annulus is horizontal (“Far=H”), vertical (“Far=V”), or empty (“No Far”). The orientation of the near-surround was ±45° (“400 ms N=45°”) or ±20° (all other conditions). The results showed a lower TI for “Far=V” compared with “Far=H” and “No Far”. (**C**) Same as panel B, for JND data. Error bars are ±1SEM.

**Fig. 6.**
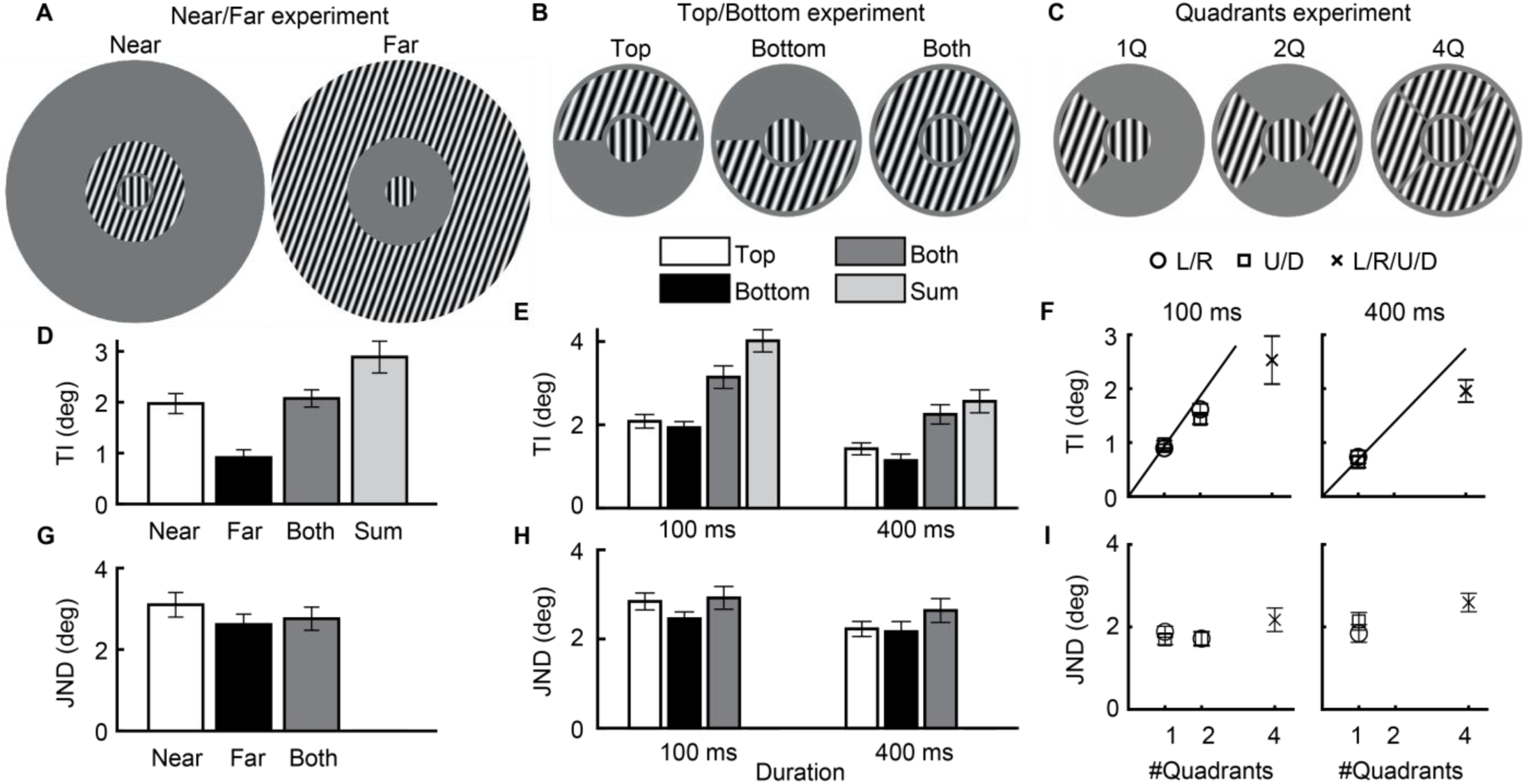
Sub-linear additivity of bias for an increased area of surround. (**A-C**) Illustrations of the used stimuli. The “Near” annulus in “Near/Far” (panel A) has a size identical to the full annulus in the “Top/Bottom” and the “Quadrants” experiments (panels B and C). The background was always uniform gray. (**D**) TI averaged across observers from component contexts (“Near” and “Far”), from the combined context (“Both”), and for the prediction based on additivity (“Sum” of the TI of the two components) (400 ms presentation duration). Sub-additivity is evident by Both<Sum. (**E**) TI for components, “Both”, and the “Sum” prediction, as in panel D. (**F**) TI averaged across observers of one quadrant surround (“1Q”: left/right/up/down), two quadrants (“2Q”: left and right, or up and down), and four quadrants (“4Q”), pooled from all “Quadrants” experiments (Table 1). The continuous line indicates the prediction from the “1Q” measurement based on additivity. The results showed, under all conditions, that an increase in the context area led to a sub-linear increase in the influence of context. (**G-I**) JND measurements. Error bars are ±1SEM.

**Fig. 8.**
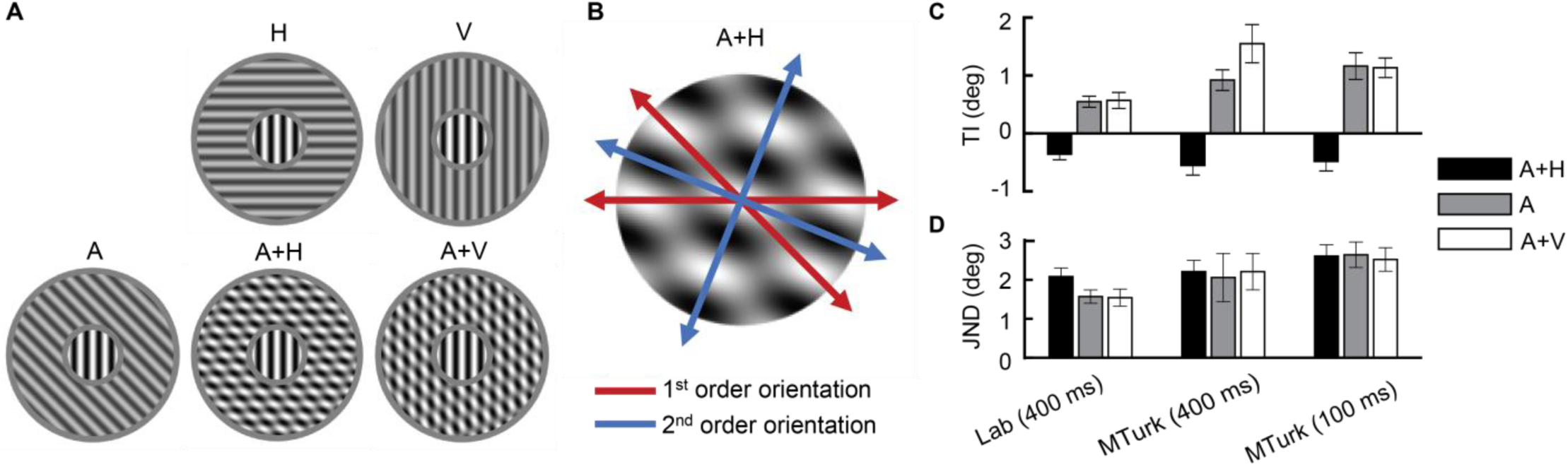
Non-linear competition between first- and second-order orientations. (**A**) Used surrounds were an oriented grating (“A”, ±20°), and mixtures of the oriented surrounds with vertical or horizontal surrounds (“A+V” and “A+H” plaids, respectively). (**B**) Illustrations of the first and second order orientations of the A+H stimuli from panel A. (**C**) Measured TI magnitudes, averaged across observers, for the different surrounds. Note that the vertical and horizontal surrounds alone do not cause TI because they are not titled relative to the near-vertical target. The results showed TI following the first-order orientation only in the absence of a higher-order oriented structure, as indicated by the measurement of negative TI under the A+H condition. (**D**) Average JNDs. Error bars are ±1SEM.

**Fig. 9.**
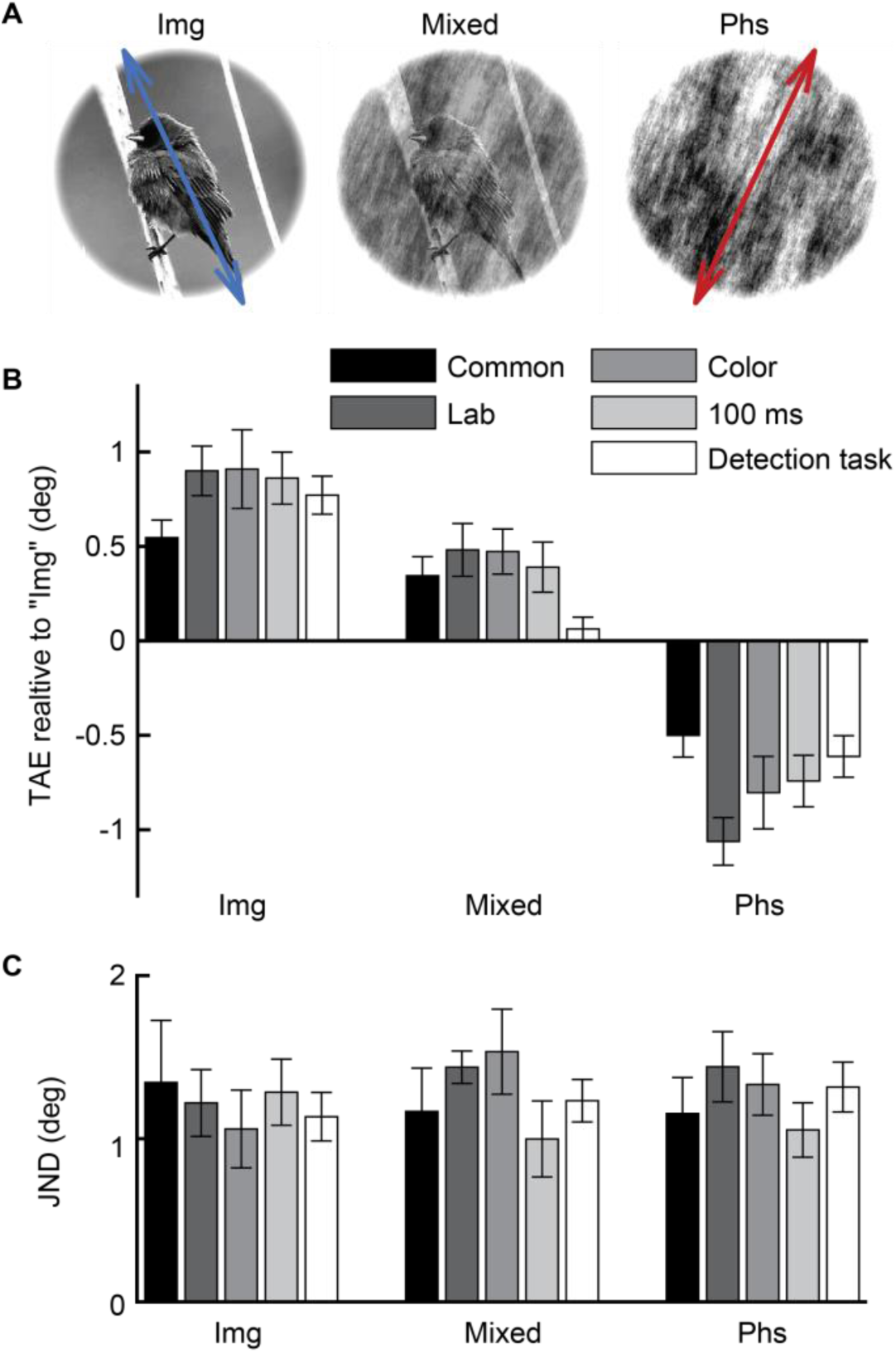
Non-linear competition between a natural image and a phase-scrambled image. **(A)** Stimuli illustrations. (**B**) TAE, averaged across observers, is shown, with the sign determined by the image content orientation (i.e., opposite the phase content orientation). In the experiments reported here, we measured the TAE due to exposure to biased images (“Img”, which are unmodified natural images with content oriented around +25° or -25° to vertical, Dekel & Sagi, 2015), phase scrambles of the biased images (“Phs”), or a mixture of the two stimulus types having opposite orientations (“Mixed”, a +25° image mixed with a phase scramble of a -25° image, or vice versa). Observers either performed an image classification task (animal vs. no animal), or a detection task (adaptor-present vs. adaptor-absent). The results showed, for the “Mixed” stimuli, a significant tilt in a direction consistent with the image content only in the presence of the classification task (not for the detection task), that is, additivity that depends on the task. The results also showed no significant change in TAE magnitude due to phase scrambling, as indicated by a similar TAE magnitude for “Img” and “Phs” (note that the sign of TAE is the same; in the plot the sign is flipped to reflect the structure of the experiment under the “Mixed” condition). (**C**) Same as panel B, for JND data. Error bars are ±1SEM.

Overall, the results suggest that in the absence of interactions during processing or selection, context-dependent biases are approximately additive (see the Discussion).

## Methods

### Observers

The experiments reported here were performed in a laboratory or with a web-based interface through the Amazon Mechanical Turk (MTurk). In the laboratory experiments, *N =* 23 observers (20 females, 3 males, aged 22-40) with normal or corrected-to-normal vision participated, and provided their written informed consent. In the MTurk experiments, *N =* 270 observers participated (reported age of 38 ± 10, Mean ± STD, median of 35, in the range 20-70; gender information was usually not collected). The web-based study was reviewed by the Institutional Review Board (IRB) of the Weizmann Institute of Science and deemed exempt from the collection of informed consent forms. The typical selection criteria for MTurk observers were as follows: ‘PercentAssignmentsApproved’ ≥ 99, ‘NumberHITsApproved’ ≥ 2000, ‘LocaleCountry’ = ‘US’. All observers were naïve to the purpose of the experiments. The work was carried out in accordance with the Code of Ethics of the World Medical Association (Declaration of Helsinki), and was approved by the Institutional Review Board (IRB) of the Weizmann Institute of Science.

### Apparatus

#### Laboratory setup

The stimuli were presented using dedicated software on a 22” HP p1230 monitor operating at 85Hz with a resolution of 1600×1200 that was gamma-corrected (linearized). The mean luminance of the display was 26.06 cd/m^2^ (TAE experiments, excluding the conditions with image adaptors) or 48.96 cd/m^2^ (TI experiments, and TAE experiment with image adaptors), in an otherwise dark environment. The monitor was viewed at a distance of 100 cm.

#### Amazon Mechanical Turk setup

To display time-accurate full-screen stimuli in arbitrary MTurk worker machines, we developed a dedicated web-browser client. The client was implemented in Javascript using WebGL, and achieved approximately single-frame accuracy on standard machines that run the Chrome, Mozilla, or Opera browsers and have a GPU-enabled WebGL implementation (this hardware support is indicated by the ‘failIfMajorPerformanceCaveat’ Javascript API flag). In addition to the browser and WebGL requirements, clients were required to use a PC rather than a tablet or a smartphone (as indicated by standard API flags), and the client machine had to pass a simple performance test (intended to ensure that the machine can achieve the required time accuracy). During an experiment, timing information estimations were collected (obtained using the ‘requestAnimationFrame’ callback). Timing diagnostics showed that most trials, of most observers, measured nearly perfect single-frame accuracy. For example, we measured for each observer the 95% worst-case presentation duration errors (across trials, relative to the observers’ average). This showed that for practically all observers (99%), the 95% worst-case error was less than 20% (e.g., setting a duration of 100 ms resulted in 95% CI [80, 120]). Similarly, inter-observer differences were negligible (*SD* of ∼5 ms). As such, we deemed the timing accuracy of the setup to be more than sufficient for the purpose of the current study. The display luminance, gamma calibration, sitting distance, and environmental lightning conditions were only controlled to the degree reflected in the task performance (see below). We note that all statistical comparisons we report are within-observer (paired), hence reflect measurements from the same display. Moreover, in all comparable experiments, we found the crowdsourcing setup to be nearly as reliable as that of the lab.

### Stimuli and tasks

All stimuli (see Fig. 2) were presented using dedicated software on a uniform gray background. To begin stimulus presentation in a trial, observers pressed the spacebar (self-initiated trials). In the laboratory, observers were instructed to fixate on the center of the display before stimulus initiation. Responses were provided using the left and right arrow keys. We measured the change in the perceived orientation of a target stimulus following exposure to an adapting oriented stimulus (TAE) or due to a surrounding oriented stimulus (TI). In MTurk, the stimuli, defined in pixels, were resized using linear interpolation to maintain a fixed proportion relative to the full-screen monitor resolution. To minimize resizing artifacts, the resizing multiplier was rounded to the nearest lower multiple of 0.25 (e.g., resizing by a multiplier of 1.25 instead of 1.33). Under all TAE conditions (and separately under all TI conditions), the stimuli used to measure the perceived orientation (targets) were nearly identical.

#### 1- and 2-back experiments (TAE)

The following presentation sequence was used (Fig. 4A): a blank screen (600 ms), 2-back Gabor adaptor (oriented -20° or +20° to vertical, 50 ms), a blank screen (600 ms), 1-back Gabor adaptor (±20°, 50 ms), a black screen (600 ms), and a near-vertical Gabor “test” (50 ms). To clarify, by *n*-back adaptor we refer to the adaptor that is *n* time frames before the test presentation (see Fig. 4A). In the “AAAAT” and “AT/AAT/AAAT/AAAAT” experiments (see Table 1), the number of adaptor presentations in the presentation sequence of a trial was always four (“AAAAT” experiment) or randomly between one and four (“AT/AAT/AAAT/AAAAT” experiment). Presentations were separated by blank screen (600 ms), similar to the two-adaptor case detailed above. All presentations were randomized independently (location and orientation). Gabor patches were 50% Michelson contrast with Gaussian envelope σ = 0.42°, spatial frequency wavelength λ = 0.3°, and random phase. Two versions of the experiment were run: fixation and periphery. In fixation, adaptors and targets were presented at the fixated center of the display, and targets were oriented -9° to +9° (in steps of 1°). In the periphery, adaptors and targets were presented either at the left or right of the fixation (at ±1.8°). The target was presented either at the same side as the adaptors (retinotopic) or at the opposite side (non-retinotopic), randomly. Targets were oriented from -12° to +12° (in steps of 2°). Observers were instructed to inspect the adaptors and the target presentations, and then to report whether the orientation of the near-vertical target was tilted clockwise or counter-clockwise relative to vertical (no feedback, except during a brief pre-experiment practice). Four peripheral crosses co-appeared with the target to facilitate discrimination between the adaptor and the target.

Note that our analyses were only for the 1- and 2-back presentations, whereas the design of some conditions permitted further conditioning of the analysis to the 3- and 4-back presentations. These data were not shown, because insufficient trials per condition due to a combinatorial explosion, which prevented drawing meaningful conclusions from the analysis. In addition, the effect from the 3- and 4-back adaptors was weaker and less consistent than that observed for the 1- and 2-back adapters.

#### Img+Phs experiments (TAE)

In these experiments, we measured the TAE caused by an adaptor presentation using a subsequently presented Gabor target (Fig. 9).

*‘Biased’ image adaptors* were unmodified natural images depicting animals and plants that were selected to maximize the orientation content around a particular combination of orientation and spatial frequency. The images were obtained using a procedure similar to that of (Dekel & Sagi, 2015). Briefly, sub-images of 256×256 pixel size were cropped from ImageNet images (Deng et al., 2009), were windowed using a circle with smoothed edges, and normalized to have an RMS contrast of 23% and an average pixel value of 127 (i.e., background). Importantly, the sub-images were selected based on the maximization of the response of an orientation filter at a specific frequency band, leading to a “biased” set of unmodified images having similarly oriented content. The orientation filter had an orientation profile of a Gaussian around the orientation 25° counter-clockwise to vertical, with a standard deviation of 15°. The frequency band had a frequency profile of a second-order Butterworth, with half-responses at 6.75 and 33.75 cycles/image. Manual pruning was used to ensure that only intelligible images of animals and plants were used. Here, the process resulted in a pool of 47 animal and 21 no-animal images (out of ∼6 million sub-image candidates, cropped with overlapping from 279,827 ImageNet images). By mirroring images, the orientation of the content could be switched from +25° to -25°. During a session, presentations of images of both orientations were randomly mixed (+25° and -25°), and each image could be presented multiple times, but always for the same orientation (randomized between observers by using a pool of 8 random assignments).

*Phase scramble adaptors* were obtained by applying the following procedure to a ‘biased’ image: applying a circular window, randomizing the phase content of the image, applying the circular window again, and then normalizing the image to have an RMS contrast of 23% and an average pixel value of 127. With color images, each RGB color channel was scrambled separately (in the standard RGB color space, which appeared to produce fewer artifacts than scrambling at an HSV or HSL color space). Another method (not used here), which preserves the phase coherence between the color channels, is to use the same phase noise in all channels (Galerne, Gousseau, & Morel, 2011).

*Mixture adaptors* were a mixture between a biased image and a phase scramble of a biased image, each at half amplitude, and each corresponding to an opposite orientation content (biased +25° and scrambled -25°, or vice versa; note that at MTurk, the contrast remained identical for both components, even if it was not at 50%).

*Gabor targets* were Gabor patches, oriented from -6° to +6° (in steps of 0.75°), with a random phase. In the laboratory setting, Gabor targets were 50% Michelson contrast with σ = 0.47° and λ = 0.33°. In the MTurk setting, Gabor targets had an amplitude of 64 gray levels (out of 256, i.e., corresponding to a contrast of 50% in a linearized display), σ = √2λ, and λ = ∼2% of screen height.

*The trial structure* used the following presentation sequence: a blank screen (350 ms), adaptor presentation (content oriented -25° or +25° to vertical; 300 or 100 ms; equal width and height, height in lab: 5.64°, height in MTurk: approx. 30% of the total monitor height), a blank screen (600 ms), and a near-vertical Gabor “target” (50 ms). A double-task was used. In the first reply, observers reported the orientation of the target (presented second). In the second reply, observers either reported the content of the image adaptor (animal/no-animal, each 50% of trials; the task showed ∼90% and ∼78% accuracy for natural and mixture adaptors, respectively), or reported whether the adaptor appeared or not (adaptor-present in 75% of trials; when the adaptor was absent, a blank screen was presented instead; the task showed ∼97% accuracy). There was no feedback in any of the tasks, except during a brief pre-experiment practice.

#### Contrast experiments (TAE)

The contrast experiments (Fig. 6) had a Gabor adaptor version and an image adaptor version. In the Gabor adaptor version, the structure of trials in the “1- and 2-back” fixation experiments was used, without the 2-back presentation and its preceding ISI (i.e., “AT”), and with varying Michelson contrast of the 1-back adaptor (11%, 33%, and 100%). In the image adaptor version, the structure of trials from the “Img+Phs” experiment was used, with only biased images (no phase scrambles), with detection as the second task in the double-task design, and with varying contrasts of the image adaptor (10%, 30%, 100%, and technically also 0% for the detection task).

#### TI experiments

Stimuli (e.g., Fig. 1A right) were presented at the center of the display, and consisted of a sine-wave circular “target” (oriented near-vertical from -9° to +9° in steps of 1°), a full, partial, or empty “near-surround” sine-wave annulus (oriented -20°, +20°, -45°, +45°, -90°, 0°, or combinations), and a full or empty “far-surround” sine-wave annulus (oriented -20°, +20°, -90°, or 0°). A partial near-surround was either a single half (top or bottom), a single quadrant (left, right, up, or down), two quadrants (left and right, or up and down), or four quadrants (left, right, up, and down with smoothed edges). Under laboratory conditions (MTurk conditions), the following sizes were used: target circle radius = 0.6° (∼3.3% of screen height), gap between target and near surround = 0.15° (∼1%), near surround radius = 1.2° (∼6.6%), gap between near and far surrounds = 0.15° (∼1%), far surround radius = 3° (∼16.6%), and spatial frequency wavelength λ = 0.3° (∼1.6%) for all sine-wave gratings. The phase was separately randomized for each stimulus component (target, near surround, quadrants of the near surround, and far surround). The sine-wave gratings had a contrast of 100%, except in the experiments with overlapping combinations (2^nd^-order stimuli, Fig. 8), where the “A”, “H”, and “V” components (−20°/+20°, -90°, and 0° orientation, respectively) each had a Michelson contrast of 50% in the lab, and an amplitude of 64 gray levels in MTurk (out of 256, i.e., corresponding to a contrast of 50% in a linearized display; note that contrast remained identical for both components even if it was not at 50%). In the experiments, observers were instructed to ignore the surrounding annuli, inspect the target (central circle), and report whether the target is tilted clockwise or counter-clockwise relative to vertical (no feedback, except during a brief pre-experiment practice). The stimuli were presented starting from 350 ms after the trial initiation, or, under a few conditions, randomly 350 to 550 ms after the trial initiation (“onset jitter”, see Table 1; this design was intended to enable a reaction time-based analysis, as in Dekel & Sagi, 2019). The stimulus was presented for 100 or 400 ms.

### Procedure

Most laboratory observers participated in multiple experiments in this study, with at least three days break between sessions (within and between experiments). All MTurk observers participated in a single experiment, for a single daily session (as indicated by having a unique ‘Worker ID’). In all experiments, we required a low rate of finger errors, achieved by persistent coaxing of laboratory observers, and pruning of MTurk observers (see below). Each daily session was preceded by a brief practice block with easy stimuli (this practice was repeated until close-to-perfect accuracy was achieved, manually in the laboratory, and automatically in the MTurk experiments). Each daily session consisted of either four or five blocks, with breaks lasting at least two minutes between blocks. The number of trials per block was determined such that the block duration was approximately five minutes (fewer trials per block when the trial duration was longer, see Table 1). When different trial types were mixed in a block, the number of trials per type was approximately equal, with the exception of the “quadrants” experiments where 2/3 of the trials were used for the single quadrant types, and 1/3 of the trials were used for the two or four quadrant types. The intended number of observers per condition was ∼10 in the lab, and ∼15 in the MTurk, which seems more than sufficient for the type of effects considered here. Under a few conditions, we doubled the number of observers following the data analysis, to ensure the validity of the found results.

### Analysis

#### Fitting perceived orientation

The magnitude of TAE and TI was calculated based on the reported orientation (clockwise vs. counterclockwise) of the near-vertical target (the Gabor patch for TAE, and central sine-wave circle for TI). First, for each context condition, a cumulative normal distribution (with lapse rates) was fitted to the psychometric function of %clockwise reports as a function of target orientation. Then, using the fitted function, the perceived vertical orientation (PV) of the target was calculated as the interpolated orientation having an equal probability for clockwise and counter-clockwise reports (50%). This was achieved with Psignifit 3.0 software (Fründ, Haenel, & Wichmann, 2011). Then, the TAE and TI magnitudes were calculated as half the shift in PV between two opposing adaptor or surround orientations (e.g., +20° vs. -20°).

To calculate JNDs (just noticeable differences), we used the slope of the fitted psychometric function. Specifically, we used the width of the interval over which the fitted cumulative Gaussian function rises from 0.25 to 0.75 (corresponding to 75% accuracy, or, put differently, to 1.35σ where σ is the fitted standard deviation). This interval was corrected so that the upper and lower lapse rates, as measured by the fit, do not affect the JND. The JNDs were calculated separately and then averaged over the -20° and +20° contexts.

When combining measurements of TAE or JND across experimental days or spatial positions (left and right, or up and down), the above calculations were performed separately for each measurement and the results were then averaged.

#### Observer and trial pruning

For the MTurk observers, the sessions were pruned, based on the parameter thresholds described in Table 2; the counts of pruned observers are presented in Table 1. It can be seen that the thresholds used were relatively lenient, and nearly none of the cooperative observers who performed the task(s) as instructed should have been pruned. Importantly, most observers were either similar to the laboratory observers in terms of measured performance, or nearly at chance level performance, so pruning, which aimed to disqualify the uncooperative observers, was usually mostly straightforward. In addition, for both MTurk and the laboratory data, trials were pruned if the orientation discrimination task RT was faster than 300 ms, or slower than 10 seconds (see Table 1 for the counts of pruned trials).

**Table 2.** Observer pruning for MTurk.

| MTurk observer pruning | “Img+Phs”<br>Orientation disc.<br>(first task) | “Img+Phs”<br>Recognition task<br>(second task) | “Img+Phs”<br>Detection task<br>(second task) | TI experiments<br>Orientation disc.<br>(single task) |
| --- | --- | --- | --- | --- |
| Mean RT > | 430 ms | 650 ms | 650 ms | 450 ms |
| Mean RT in the fastest block > | 430 ms | 450 ms | 450 ms | 450 ms |
| Mean accuracy > | 65% | 60% | 90% | 65% |
| Mean accuracy in the worst block > | 65% | 60% | 85% | 65% |
| Reduction in the mean accuracy<br>from the best to the worst block < | 0.2 | 0.2 | 0.2 | 0.2 |
| Reply heterogeneity < | 12% with Det.<br>20% with Rec. | - | - | 17% for Q1/Q2<br>25% otherwise |

#### Statistics

All relevant statistical tests were two-tailed. All *t*-tests were paired (i.e., repeated-measures).

### Modeling

#### Additive model

In the TAE experiment, we found additivity (see the Results). To model this, we describe each adaptor’s influence by a scalar indicating the weight of its influence; the sign of the weight is the sign of the adaptor (clockwise, counterclockwise, and absent, indicated by +, -, and 0, respectively), and the predicted PV is the sum. This can be written as:

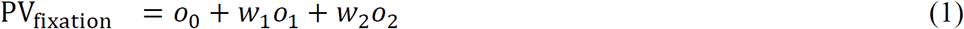

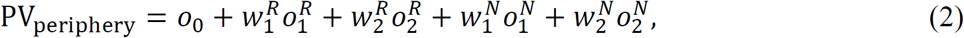

where PV is the perceived orientation, *o*_0_ is the reference (unadapted) orientation, *o*_*i*_ is the *i*-back adaptor orientation (*R* and *N* indicate the retinotopic vs. non-retinotopic adaptor, respectively), and *w*_*i*_ is the weight of the *i*-back adaptor.

#### Cue combination model

To interpret biases in the perceived orientation, we considered the idea that a reference frame is updated based on using the contextual orientations as cues. We modelled this idea as a Bayesian cue combination of independent Gaussian cues (Oruç, Maloney, & Landy, 2003):

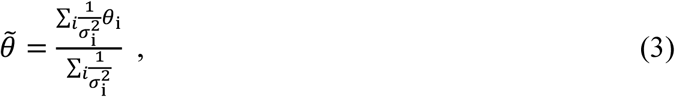

where *θ*_*i*_ and 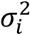 are the means and variances of the Gaussians, and 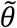 is the posterior mean. The quantity 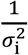 is often referred to as *reliability*. The model described each adaptor, as well as the unadapted vertical reference, act as independent Gaussian cues (Fig. 3). Specifically, the unadapted reference cue had an unknown orientation, *θ*_0_, each adaptor cue had a known orientation, *θ*_i_, and the relative reliability between each adaptor and the unadapted reference, 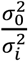, was also unknown.

To model the fixation experiment using a cue combination, let *θ*_0_ be the initial (unadapted) internal reference orientation, *θ*_1_ be the 1-back adaptor orientation, and *θ*_2_ be the 2-back adaptor orientation. The updated internal reference, inferred using Bayes’ rule and by assuming that the cues are Gaussian with variances 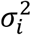, is (see Eq. (3)):

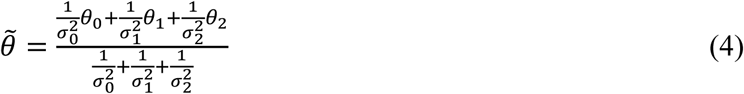

The predicted vertical orientation is minus (opposite) the change in the internal reference, which is, 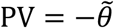. Rearranging, we get:

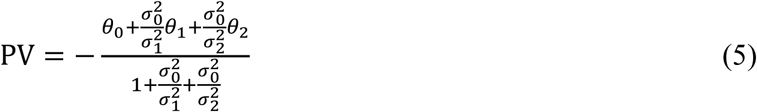

When an *i*-back adaptor is missing, we can set *σ*_*i*_ = ∞, which removes the relevant terms from the equation. Of the model parameters, the adaptor orientations are known and equal to ±20°; therefore, three unknown parameters remain: *θ*_0_ (the initial internal reference), 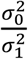, and 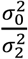.

Similarly, the equation for the periphery experiment can be obtained (five unknown parameters). Note that in both cases, the number of parameters is the same as in an additive model (Eqs. (1) and (2)). When fitting behavior to the model, nearly identical predictions were found when fitting the average observer, compared to when fitting each observer separately, and then averaging. We therefore used the latter, which is the stricter alternative in the sense that it can be more non-additive.

Of note, based on the above cue-combination model equations, the TAE, which is the change in the perceived orientation, can be derived for the case of a single adaptor as follows (TI is calculated similarly):

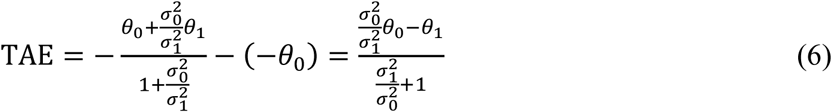

## Results

### Linear additivity for temporally separated contexts

First, we considered the change in the perceived orientation caused by temporal combinations of spatially isolated Gabor patches. Specifically, we considered two possible presentation times, 650 and 1300 ms, before a target, which we referred to as 1-back and 2-back, respectively. In each time frame, the fixated center of the display presented a Gabor tilted +20° clockwise to vertical, -20° to vertical, or was empty (Fig. 4A). We measured the change in the perceived vertical orientation of the target stimulus due to the adaptor(s)’ exposure (tilt aftereffect, TAE), and checked whether the combined influence of both adaptors can be explained as a linear sum of the isolated adaptor influences. The results showed standard TAE magnitudes of around ∼1° (Fig. 4D), and importantly, the additivity of the 1- and 2-back adaptation intervals (Fig. 4B). We analyzed this using a 3×3 repeated measures ANOVA of {-20°,+20°,empty}×{-20°,+20°,empty}, which showed the significant influence of the 1- and the 2-back presentations, and importantly, no interaction, indicating additivity (1-back: *F*_(2,16)_ = 26.99, *p* < 0.0001; 2-back: *F*_(2,16)_ = 20.25, *p* < 0.0001, interaction: *F*_(4,32)_ = 1.08, *p* = 0.38, *N =* 9 observers, Fig. 4B). To strengthen this result, we also considered a linear mixed-effects regression analysis, which showed the same findings (1-back: *t*_(77)_ = 8.81, *p* < 0.0001; 2-back: *t*_(77)_ = 4.64, *p* < 0.0001; interaction: *t*_(77)_ = -0.05, *p* = 0.95).

Similarly, using presentations in the near-periphery (±2° of visual angle, Fig. 4A), we considered the 1- and 2-back adaptors, each presented at the same side as the target (retinotopic, “R”) or at the contra-lateral side (non-retinotopic, “N”). We analyzed the data from two experiments, with four adaptor presentations before the target (AAAAT, *N =* 5 observers, *n* = 13,271 trials), or with up to four adaptor presentations before the target (AT, AAT, AAAT, or AAAAT, *N =* 6 observers, *n* = 15,608 trials). The results showed retinotopic TAEs of about 1°, non-retinotopic TAEs of about 0.25° (Fig. 4D), and importantly, additivity of the 1- and 2-back adaptors (Fig. 4C). Specifically, we applied a 4×4 repeated measures ANOVA of {R-20°,N-20°,N+20°,R+20°}×{R-20°,N-20°,N+20°,R+20°}, which showed significant 1- and 2-back adaptor influences, but non-significant interaction, for “AAAAT” (1-back: *F*_(3,12)_ = 28.71, *p* < 0.0001, 2-back: *F*_(3,12)_ = 23.98, *p* < 0.0001, interaction: *F*_(9,36)_ = 0.41, *p* = 0.92), and similarly for “AT, AAT, AAAT, AAAAT” (1-back: *F*_(3,15)_ = 3.38, *p* = 0.05, 2-back: *F*_(3,15)_ = 7.25, *p* = 0.003, interaction: *F*_(9,45)_ = 0.25, *p* = 0.98). We therefore merged the data of the two experiments (1-back: *F*_(3,30)_ = 14.96, *p* < 0.0001, 2-back: *F*_(3,30)_ = 22.81, *p* < 0.0001, interaction: *F*_(9,90)_ = 0.19, *p* = 0.99; *N* = 11 total observers from both experiments, Fig. 4C). Therefore, the results showed an additivity of peripheral adaptation, and moreover, an additivity of retinotopic with a non-retinotopic adaptation. The 3- and 4-back adapters led to a TAE (∼0.25° retinotopically); however, the effect was weaker and less consistent than that observed for the 1- and 2-back adapters; therefore, analyzing the additivity was not immediately possible (see the Methods).

Overall, across all conditions, the perceived vertical orientation (PV), following a sequence of temporally separated adaptors, can accurately be described as the sum of the isolated adaptor influences (see the Methods, Eqs. (1) and (2)). That is, the combined effect was well explained as the sum of the component influences, with non-significant deviations from additivity. One possibility of interpreting these findings is by using the theory that a bias in the perceived orientation reflects an update of an internal reference frame by using the contextual orientations as cues (Andrews, 1964; C. J. Dakin & Rosenberg, 2018). For example, an adaptor oriented +20° might shift the vertical reference from 0° to +1°, in which case a physical 0° orientation will be perceived as -1° (relative to the reference). We modelled this idea as a Bayesian cue combination of independent Gaussian cues (see the Methods; Oruç, Maloney, & Landy, 2003). Remarkably, model fitting showed that in the relevant parameter range 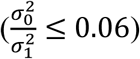, the model is practically equivalent to additivity. That is, the cue-combination predictions (in orange) were superimposed on the additive predictions (in black; see Fig. 4). Quantitatively, for both additive and cue-combination models, the resulting *R*^2^ values (the prediction errors in Fig. 4BC) showed *R*^2^ = 0.97 in the fixation experiment, and *R*^2^ = 0.99 in the periphery experiment. Therefore, the independent cue-combination model accounts for the observation of the additivity of temporally separated contexts. Note that the cue combination account provides a theory, whereas additivity is an arbitrary rule (see the Discussion). The orientation sensitivity in the tasks (just noticeable differences, JND) was measured using the inverse slope of the fitted psychometric functions (see the Methods). Results (Fig. 4EF) showed inconsistent and mostly non-significant modulation of JND by the 1-back, the 2-back, and the interaction of the 1- and 2-back adaptors. Statistics were assessed by applying the ANOVA tests described above, for fixation (Fig. 4E; 1-back: *F*_(2,16)_ = 0.30, *p* = 0.7; 2-back: *F*_(2,16)_ = 0.04, *p* = 0.9, interaction: *F*_(4,32)_ = 2.80, *p* = 0.04), and periphery (Fig. 4F; 1-back: *F*_(3,30)_ = 0.81, *p* = 0.5, 2-back: *F*_(3,30)_ = 0.06, *p* = 0.98, interaction: *F*_(9,90)_ = 0.99, *p* = 0.56; *N* = 11 observers).

### Sub-linearity for increased adaptor contrast

Next, we considered an increase in an adaptor’s contrast (Fig. 5). That is, we measured the TAE at different adaptor contrasts, and compared the result to what is expected from linearity of the TAE vs. contrast function. The results showed a lack of linearity (Fig. 5), consistent with previous reports (Blake, Tadin, Sobel, Raissian, & Chong, 2006). For example, an increase of contrast by a factor of almost 10, from 11% to 100%, led to an increase in TAE by a factor of only ∼2.5. More generally, the 100% contrast measurement was lower than the multiplication of a lower-contrast measurement by the factor of the contrast change, for Gabor adaptors (compared to 33% contrast: *t*_(8)_ = 3.42, *p* = 0.01; compared to 11% contrast: *t*_(8)_ = 2.20, *p* = 0.06; two-tailed repeated measures t-test, not corrected for multiple comparisons; *N* = 9 observers), and for ‘biased’ image adaptors (compared to 30% contrast: *t*_(9)_ = -3.50, *p* < 0.01; compared to 10% contrast: *t*_(9)_ = -3.46, *p* < 0.01; *N* = 10 observers, using the ‘Detection task’ double-task design, see the Methods). Therefore, an increase in contrast led to a sub-linear increase in TAE.

Importantly, the same Gabor stimuli were used in the Gabor contrast experiments and in the temporal separation experiments (Fig. 4BC and Fig. 5, “Gabor”). Moreover, the magnitudes of TAE were comparable. Therefore, the finding of sub-linearity here rules out the possibility that the additivity of the temporally separated contexts found above is due to measuring a TAE with a small magnitude. (It is also interesting to note that the manipulation of contrast is, in a sense, the opposite of temporal separation: sufficiently adjacent presentations, within a period of less than ∼50 ms can be assumed to be temporally integrated, and thus to be almost interchangeable with a change in the contrast of a single presentation, see, e.g., Watson & Nachmias, 1977.)

The target JND was inconsistently modulated by the adaptor contrast (Fig. 5B). Statistically, a linear mixed-effects regression analysis showed that JND as a function of contrast did not have a slope term that is significantly different from zero (Gabor: *t*_(25)_ = -1.35, *p* = 0.18; Image: *t*_(38)_ = 1.67, *p* = 0.1).

### Sub-additivity of similar, adjacent contexts

Here, we consider combinations of similar contexts (having the same grating pattern) that are adjacent in space. Contexts were presented at the same time as the target (TI). The corresponding experiment for TAE, of combining contexts presented at the same time and tested at a later time, measured magnitudes of non-retinotopic TAEs that were too small to investigate additivity (data not shown).

First, we consider the additivity of an increase in the context area (Fig. 6). We measured the TI due to a surrounding annulus that is close to the central target (“Near”), far from the central target (“Far”), or when both annuli were presented (“Both”) (Fig. 6A). Remarkably, the results showed the same TI magnitude of 2° when the Near annulus was presented either with or without the Far annulus, despite a TI of 1° caused by the Far surround alone (Fig. 6D). Indeed, comparing “Both” to the sum of “Near” and “Far” showed a large and significant difference indicating a strong sub-additivity (*t*_(20)_ = 3.77, *p* = 0.001, *N* = 21 MTurk observers, using a presentation duration of 400 ms).

Using a similar design, we measured the TI from the upper half (“Top”), lower half (“Bottom”), or full (“Both”) surrounding annulus (Fig. 6B). The results (Fig. 6E) showed for both surround halves a TI of between half to two-thirds of the TI of the full annulus, i.e., close to additivity but with a statistically significant trend toward sub-additivity (comparing “Both” to the sum “Top” + “Bottom” showed, 100 ms presentation: *p* = 0.0002, *t*_(18)_ = 4.69, *N* = 19 MTurk observers; 400 ms presentation: *p* = 0.05, *t*_(25)_ = 2.05, *N* = 26 MTurk observers).

A similar sub-additivity was found when considering different quadrants of the surrounding annulus (Fig. 6CF). For example, with a 100 ms presentation duration, the results showed a TI of ∼1° when one quadrant was presented (“1Q”: left, right, up, or down), a TI of ∼1.5° when two quadrants were presented (“2Q”: left and right, or up and down), and a TI of only ∼2.5° when all four quadrants were presented (“4Q”). That is, a strong sub-additivity is evident by the “4Q” measurement showing much less TI than four times the “1Q” measurement (see Fig. 6F) (100 ms: *t*_(9)_ = -3.71, *p* = 0.005, *N* = 10 MTurk observers; 400 ms: *t*_(16)_ = -3.22, *p* = 0.005, *N* = 17). Moreover, a weak sub-additivity is evident by the “2Q” measurement showing slightly less TI than twice the “1Q” measurement (100 ms presentation duration: *p* = 0.01, *t*_(32)_ = -2.66, *N* = 33 MTurk observers; data were pooled from a condition having an onset jitter, see Table 1; no 400 ms condition). Importantly, comparing the left/right to the up/down measurements showed that the left and right quadrants were nearly additive (“1Q” averaged 0.86°, “2Q” averaged 1.62°, and 2·”1Q” vs. “2Q” showed: *t*_(32)_ = -0.72, *p* = 0.9 corrected), whereas the up and down quadrants measured a strong sub-additivity (“1Q” averaged 0.99°, “2Q” averaged 1.44°, and 2·”1Q” vs. “2Q” showed: *t*_(32)_ = -3.53, *p* = 0.003; Bonferroni correction for two multiple comparisons). This anisotropy can possibly be explained by the fact that the sub-additivity is caused by low-level, intra-hemisphere interactions (see the Discussion).

Overall, increasing the surround area led to a sub-linear increase in TI (Fig. 6). This finding qualitatively resembles the sub-linearity observed for increased adaptor contrast (Fig. 5 and Blake, Tadin, Sobel, Raissian, & Chong, 2006). The finding in the first section, of linear additivity for temporally separated contexts, suggests that the sub-linearity observed here (Figs. 5 and 6) is caused by non-linearity in the processing of the context (see the Discussion).

Concerning target JNDs, the modulations by the surrounds were small and barely significant (∼15% changes, Fig. 6GHI). Specifically, in the “Near/Far” experiment, a repeated-measures ANOVA of {Near,Far,Both} was not significant (*F*_(2,40)_ = 2.97, *p* = 0.06; Fig. 6G). In the “Top/Bottom” experiment, the average of the “Top” and “Bottom” measurements was compared to the “Both” measurement, which showed a significant difference in the 400 ms version (*t*_(25)_ = -3.11, *p* = 0.005), but not in the 100 ms version (*t*_(18)_ = -1.59, *p* = 0.13) (Fig. 6H). In the “Quadrants” experiment, the difference between the “4Q” measurement to the averaged “1Q” measurement was not significant in the 100 ms version (*t*_(9)_ = 1.44, *p* = 0.18), and marginally significant in the 400 ms version (*t*_(16)_ = 2.53, *p* = 0.02) (Fig. 6I). Overall, changes in TI (Fig. 6DEF) were dissociated from changes in JND (Fig. 6GHI), though there might be a weak trend of increased JND alongside increased TI.

### Non-linear competition between adjacent, dissimilar contexts

A possible account of the sub-linearity observed above for overlapping or adjacent contexts is that it reflects self-inhibition (for overlapping contexts) or lateral inhibition (for adjacent contexts). Here, we considered experiments using combinations of dissimilar contexts (e.g., one tilted and one not). These experiments showed a form of competition between contexts that cannot be readily explained by self- or lateral inhibition.

First, we measured how the TI, caused by the orientation of a surrounding annulus (“near-surround”, tilted ±20° to vertical), is influenced by a far-surround annulus that is either horizontal or vertical (Fig. 7A). The results (Fig. 7B) showed about a 30% reduction in TI in the presence of vertical compared with horizontal far surround (100 ms presentation duration: *N =* 9 MTurk observers, *t*_(8)_ = -4.77, *p* = 0.002, two-tailed repeated-measures t-tests; 400 ms presentation duration: *N =* 14 MTurk observers, *t*_(13)_ = - 4.62, *p* = 0.0005). The observed effect was robust, as evident for all observers except one (22 out of 23 showing V<H), which is especially striking considering the inherent variability of the display parameters in the web-based platform.

Moreover, the interaction with the far surround was selective to the vertical far surround orientation (“H/V/E” conditions, Fig. 7B). That is, the TI for absent vs. horizontal far surrounds was almost identical as indicated by the non-significance of the null hypothesis (“No Far” vs. “Far=H”, 100 ms: *t*_(25)_ = -0.3, *p* = 1 corrected, *N =* 26 MTurk observers; 400 ms with onset jitter: *t*_(14)_ = -1.16, *p* = 0.79 corrected, *N =* 15 MTurk observers; Bonferroni correction for three multiple comparisons), whereas TI in the presence of a vertical far surround was reduced by about 30% relative to both the absent and horizontal far surrounds (“Far=V” vs. “No Far”, 100 ms: *t*_(25)_ = -3.32, *p* = 0.008 corrected, 400 ms: *t*_(14)_ = -3.53, *p* = 0.01 corrected; “Far=V” vs. “Far=H”, 100 ms: *t*_(25)_ = -5.29, *p* < 0.0001 corrected, 400 ms: *t*_(14)_ = -4.09, *p* = 0.003 corrected). It is reassuring to note the comparable TI magnitudes for identical stimuli under the different conditions: “No Far” condition here, the “Near” condition of Fig. 6AD, and the “Both” condition of Fig. 6BE (∼2° on average for the 400 ms measurement).

Next, to address the possibility that the effect is mediated by an interaction between near and far surrounds (e.g., lateral inhibition, Clifford, 2014; O’Toole & Wenderoth, 1977), we used a near surround tilted at ±45°, with the far surround oriented horizontally or vertically as before. In this experiment, the orientation difference between the near and the far surround is always the same (45° for both vertical and horizontal far surrounds), equating the interaction between the near and the far surrounds. Importantly, the results again showed a robust reduction in TI in the presence of a vertical compared with a horizontal far surround (Fig. 7B, “400 ms N=45°” condition, *N =* 12 MTurk observers of whom 11 show the effect, *t*_(11)_ = -4.45, *p* = 0.001; a presentation duration of 400 ms was used). This suggests that the effect is mediated by an interaction between the far surround and the target, rather than an interaction between the far surround and the near surround. Note that although the far surround affects the orientation of the near surround (i.e., a TI of near surround toward or away from the orientation of the far surround), this effect is probably not too large (∼±2°); hence, it seems unlikely to lead to a large change in the TI magnitude (i.e., TI from 43° vs. 47° seems unlikely to lead to a ∼25% reduction in the effect magnitude). Overall, the results showed that the interaction between the target and the near surround (as measured by TI) interacts with the interaction between the target and the far surround (a three-way interaction: target × near surround × far surround). Put differently, the results showed that competition exists between the different contextual interactions of the target (see the Discussion). A possible interpretation of the results is given by considering the figure-ground segmentation of the stimulus, which may depend on the far-surround orientation (von der Heydt, Macuda, & Qiu, 2005) (see the Discussion).

To address the possibility that the reduction in TI is mediated by an improved orientation acuity in the presence of a vertical orientation cue in the far surround (Wei & Stocker, 2017), we compared JNDs between vertical and horizontal far surrounds (Fig. 7C). JNDs were calculated as the inverse of the slope of the psychometric function (i.e., percent clockwise reports as a function of target orientation, see the Methods). The results showed minor and non-significant modulated in JND in the presence of the vertical compared with the horizontal far surround, and even a nonsignificant deterioration in three of the five conditions (“100 ms”: *t*_(8)_ = -0.43, *p* = 0.67; “400 ms”: *t*_(13)_ = 1.61, *p* = 0.13; “100 ms H/V/E “: *t*_(25)_ = -0.94, *p* = 0.35, “400 ms H/V/E”: *t*_(14)_ = -0.19, *p* = 0.84; “400 ms N=45°”: *t*_(11)_ = 0.65, *p* = 0.52). Therefore, the main effect was not explained by improved orientation acuity.

### Non-linear competition between overlapping, dissimilar contexts

Next, we considered combinations of overlapping contexts that are dissimilar (Figs. 8 and 9). These experiments are important, because natural stimuli are composed of dissimilar orientations. Moreover, the findings may prove diagnostic for comparing the different theoretical accounts of bias (see the Discussion).

We first considered the TI caused by a tilted surround (“A”), a mix of a tilted and a vertical surround (“A+V” plaid), and a mix of a tilted and a horizontal surround (“A+H” plaid) (Fig. 8A). Importantly, in terms of first-order orientation (i.e., contrast), the mixtures maintain the orientation of “A”, with the added vertical or horizontal orientations having no tilt relative to the vertical target. However, unlike a prediction based on the first-order orientation, the results (Fig. 8C) showed that TI depends on the identity of the mixed grating, as indicated by a repeated measures ANOVA (Lab, 400 ms: *F*_(2,18)_ = 27.76, *p* < 0.0001, *N =* 10 observers; MTurk, 400 ms: *F*_(2,12)_ = 23.95, *p* = 0.0001, *N =* 7 MTurk observers; MTurk, 100 ms: *F*_(2,22)_ = 22.17, *p* < 0.0001, *N =* 12 MTurk observers). Specifically, the “A+V” and “A” conditions measured a similar TI (differences showing *p* > 0.05 in each of the three experiments), whereas the “A+H” condition measured much less TI than “A+V”, and much less TI than “A” (all post-hoc *t*-tests showing *p* < 0.005, Bonferroni corrected for three multiple comparisons). Moreover, the “A+H” measurement was typically negative (*p* < 0.05 in each of the three experiments, again corrected for three multiple comparisons). This finding shows the non-additivity of the mixing.

The direction of TI can be explained by the perceived, second-order orientation (Dakin, Williams, & Hess, 1999; Smith, Clifford, & Wenderoth, 2001; Wei, Zhou, & Chen, 2018). As evident in a visual inspection (Fig. 8B), the checkerboard pattern is grouped along either the orientation of the elements, or the orthogonal orientation. For “A+V” with the near-surround titled -45°, this results in either a -22.5° or a +67.5° second-order orientation. For “A+H” with the near-surround titled -45°, this results in either a - 67.5° or a +22.5° second-order orientation. (We quantitatively verified this claim by finding the orientations with the most Fourier energy in the “contrast” of the stimuli, where “contrast” was calculated as the squared difference between the stimuli image and the background value.) Importantly, the sign of TI depends on the orientation difference between the surround and the target: at ∼5 to ∼65° orientation differences, the shift in the perceived orientation is in the opposite direction relative to the surround orientation (“repulsive effect”), whereas orientation differences of ∼80° lead to a shift of the perceived orientation in the direction of the surround (“attractive effect”, also known as the indirect tilt illusion) (“The tilt illusion: Phenomenology and functional implications,” 2014). Therefore, under the “A+V” condition, the first-order orientation content (e.g., repulsive from -45°) leads to the same sign of TI as the second-order orientation (repulsive from -22.5°, or attractive toward +67.5°). However, under the “A+H” condition, the first-order orientation content (e.g., repulsive from -45°) leads to an opposite sign of TI than does the second-order orientation (attractive toward -67.5°, or repulsive from +22.5°). Therefore, under the mixed conditions, the direction of TI was consistent with the second-order (or perceived) orientation. This account suggests that when first- and second-order orientations are not consistent, the influence follows the higher-order orientation.

It is important to emphasize that although both adjacent (Fig. 7) and overlapping (Fig. 8) combinations with vertical and horizontal gratings showed non-linearity, the measured direction of influence was opposite (V>H for overlapping, and H>V for adjacent, see Fig. 7 and Fig. 8, respectively; see the Discussion). This observation is particularly interesting when considered in the context of phase scrambling approaches, showing that the phase (i.e., relative position) of an oriented component determines its influence.

Modulations of JND by the surround were perhaps evident in the lab data (*F*_(2,18)_ = 5.78, *p* = 0.01), with “A+H” showing worse sensitivity (higher JND) (Fig. 8D). However, this trend was entirely absent from the MTurk data (400 ms: F_(2,12)_ = 0.06, *p* = 0.94; 100 ms: F_(2,22)_ = 0.13, *p* = 0.87; Fig. 8D).

### Non-linear competition between natural and phase-scrambled images

Finally, we considered combinations of overlapping contexts that are natural vs. reduced. To this end, we used ‘biased’ image adaptors, which are unmodified natural images that were selected to maximize the orientation content around a particular orientation (Dekel & Sagi, 2015). Observers were exposed to biased images (“Img”, with content oriented around +25° or -25° to vertical), phase scrambles of the biased images (“Phs”, the same oriented content as “Img”, but with scrambled high-order image content), and a mixture of the two image types (“Mixed”). Importantly, the “Img” and “Phs” components of the mix were always taken from opposite orientation groups (−25° with +25°, or vice versa). Therefore, TAE under the mixed condition could either reflect the image component, the phase scramble component, or neither (additive).

The results showed that the TAE under the “Mixed” condition was in a direction consistent with the image content, that is, non-additive (Fig. 9). To quantify this effect, we compared the measurement under the “Mixed” condition to the average of the “Img” and “Phs” conditions, and found a significant difference (*N =* 9 MTurk observers, *t*_(8)_ = 2.39, *p* = 0.05 two-tailed repeated measures *t*-test, Fig. 9, “Common”). This result was replicated when tested under laboratory conditions instead of using the Mechanical Turk (*N =* 10 observers, *t*_(9)_ = 4.24, *p* = 0.002), when using color images instead of grayscale (*N =* 10 MTurk observers, *t*_(9)_ = 3.28, *p* = 0.01), and when using a briefer image adaptor presentation duration of 100 ms instead of 300 ms, to rule out the influence of eye movements (*N =* 11 MTurk observers, *t*_(10)_ = 3.17, *p* = 0.01).

Interestingly, when a detection task was used instead of the image-content classification task, the results showed additivity, i.e., the “Mixed” measured the average of “Img” and “Phs” (as indicated by the non-significance of the null hypothesis, *N =* 19 MTurk observers, *t*_(18)_ = 0.28, *p* = 0.78). The absence of non-linearity with the detection task suggests that the image classification task is implicated in mediating the competition between the contexts (see the Discussion). Moreover, this finding rules out the possibility that the direction of TAE in the “Mixed” condition reflects a higher local contrast in the “Img” stimuli (since phase scrambling maintains the global but not necessarily the local contrast in the image; see also Stojanoski & Cusack, 2014). (We note that the “Img” and “Phs” components indeed measured a minor and practically negligible difference, with a slightly higher “Img” TAE in most conditions, and a slightly above-zero “Mixed” TAE for the “Detection task”, *p* = 0.04, see Fig. 9.) Overall, the results suggest that selection between the component influences is task dependent. Specifically, when the task required attending to the image component over the phase-scrambled component, the TAE followed the image.

The JNDs of the target orientation discrimination task were mostly independent of the adaptor type (see Fig. 9C). Statistically, a repeated-measures ANOVA of {Img,Phs,Mixed} showed that changes in JND were not significant in all experiments (*p* > 0.4), with the exception of the “Color” version that was marginally significant (F_(2,18)_ = 4.22, *p* = 0.03), possibly a statistical coincidence.

## Discussion

We measured bias in the perceived orientation due to different spatiotemporal combinations of contexts. Our investigation focused on meaningful qualitative patterns, which we believe to be general, because it was impractical to systematically test all combinatorial combinations of all context types.

### Additively of separated contexts

We found that for adaptors that were presented at separate times (600 ms ISI), the combined influence was explained as an additive sum of the component effects (Fig. 4, see Eqs. (1) and (2)). Notably, there was no evidence for sub-additivity (as indicated by linearity for orientation repetition) and no evidence for selection (as indicated by linearity for orientation switch and side repetition). These findings are strengthened by a previous report on additivity of the TAE from adaptors and from learned expectations (Pinchuk-Yacobi, Dekel, et al., 2016). Further, Magnussen and Kurtenbach (1980) found additivity of the TAE and the TI. Overall, it seems that contexts that are sufficiently separated in time interact additively in the measured influence, unlike commonly held beliefs.

An important observation concerning the additivity is that it refers to a specific measurement scale of orientation: degrees. Using a measurement scale that is not linear with degrees in the tested range (∼2° change in the perceived orientation) would not show an additivity of measurements. For example, if the orientation changes were represented as the logarithm of the orientation difference, then additivity would be evident for the exponent of the measurements.

Theoretically, we expect additivity in degrees from an optimal calibration of the reference orientation (see the cue combination theory below), and from population vector readout theories (whereby sufficiently small context-dependent changes in the representation are linear with the read-out bias in degrees) (Georgopoulos, Schwartz, & Kettner, 1986; Gilbert & Wiesel, 1990; Schwartz, Sejnowski, & Dayan, 2009). Our findings are in line with the existence of multiple, separate mechanisms of adaptation (Bao & Engel, 2012; Gekas, McDermott, & Mamassian, 2019), and it remains to be seen whether additivity is only evident when aggregating over separate mechanisms, or also within a mechanism (as long as the mechanism is not saturated).

### Non-additivity of overlapping or adjacent contexts (TI and TAE)

When the contexts were adjacent or overlapping in space, the combined influence could not be generally explained as an additive sum of separate influences (i.e., a non-linear interaction). Here, we parsed the absence of additivity using two terms: sub-additivity, when a combined influence is lower than the sum of the separate influences, and selection, when the combined influence reflects more one of the components than the others.

#### Sub-additivity

For increased context strength, we found sub-additivity. That is, an increase of the context contrast (Fig. 5) or area (Fig. 6), which leads to a sub-linear increase in bias. A sub-linear increase in bias for increased contrast was observed previously for the TAE (Blake et al., 2006; Harris & Calvert, 1989), but here, the results can be interpreted in light of the finding of additivity (Fig. 4; using the same experimental design). Moreover, an increase in adaptor presentation duration (which is arguably similar to a change in contrast, Watson & Nachmias, 1977), is also known to lead to a sub-linear increase in TAE (Dekel & Sagi, 2015; Sekuler & Littlejohn, 1974). Additivity for increased context area was, to the best of our knowledge, never directly tested. (An experiment by Song & Rees, 2018 is similar to the Top/Bottom experiment in Fig. 6B, but in their design the central target circle was halved along with the surround, so it is unclear whether the findings reflect the change in the target or in the surround. Still, their results of sub-additivity are consistent with our own.) Overall, the reports of sub-additivity seem ubiquitous. Moreover, electrophysiological investigations of contextual influence seem consistent with sub-linearity, though the mapping from neural activations to perception is not necessarily straightforward (see, e.g., the results reviewed by Carandini & Heeger, 2012).

Interestingly, the results measured a spatial anisotropy of the additivity. Specifically, left and right combinations were nearly additive, whereas up and down combinations measured a strong sub-additivity (Fig. 6C). The anisotropy in additivity was not explained by anisotropy of TI of isolated components: the left, right, up, and down quadrants measured nearly identical TI. A possible interpretation of this finding is that the sub-additivity of combinations reflects low-level, intra-hemisphere interactions. This claim seems consistent with a suggestion by Song & Rees (2018), based on fMRI analysis, that TI reflects intra-hemisphere integration. Alternatively, the interaction between the components may depend on the configuration (parallel vs. collinear).

#### Selection

For combinations of dissimilar contexts (Figs. 7-9), we often found a selection between the components, whereby the higher-level or task-relevant context was selected over a lower-level or a task-irrelevant context.

We first considered the case where a “neutral” context is adjacent to a tilted context. By neutral, we refer to vertical or horizontal orientations, which cannot cause TI in the vertical target (due to symmetry). The results showed reduced or unchanged TI. Specifically, the TI from a tilted near-surround context was reduced in the presence of a vertical, but not horizontal, far surround context (Fig. 7). Moreover, the reduction in TI cannot be explained by an interaction between the near and the far surrounds. Specifically, under a condition with a fixed orientation difference between the near and the far surrounds, we still observed a change in TI due to the far surround orientation (Fig. 7, “N=45°” condition). Therefore, the results indicate an interaction between the far surround and the target in mediating the effect. This finding may be interpreted as a consequence of figure-ground segregation. Specifically, when both far surround and target are vertical, the near surround may appear to “occlude” a vertical background (inspect Fig. 7A). The interaction between two differently segmented regions may be reduced, as measured by lower TI (Qiu, Kersten, & Olman, 2013; von der Heydt et al., 2005). Still, we note that here the target, near, and far surrounds were separated by gray rings, to reduce such effects.

When a neutral and a tilted context were mixed in the same position (i.e., overlapping), the second-order orientations of the resulting plaid pattern determined the sign of the TI (Fig. 8). Specifically, for TI caused by a 45° tilt, a mix with a horizontal orientation inverted the sign of the TI, whereas a mix with the vertical did not significantly change the TI. Given the results with the adjacent contexts above, it is clear that the TI is determined by the arrangement of the contexts’ orientations: H<V when overlapping (Fig. 8), and V<H or V=H when adjacent (Fig. 7, “400 ms N=45°” condition). This observation is particularly relevant for phase scrambling approaches, showing that the phase (i.e., relative position) of an oriented component determines its influence. That is, the TI cannot be reduced to basic (first-order) orientation processing circuits. In the current case, the direction of TI could be described by the second-order orientation (see Dakin, Williams, & Hess, 1999; Smith, Clifford, & Wenderoth, 2001; Wei, Zhou, & Chen, 2018). Possibly, in competition between first and second-order orientations, the TI follows the higher order orientation, consistent with the Gestalt approach to visual perception (Herzog, Thunell, & Ögmen, 2016; Jäkel, Singh, Wichmann, & Herzog, 2016). (An alternative account is that both first and second order orientations receive a fixed weight, but that the “direct effect” from the second-order orientation is stronger than the “indirect effect” from the first-order orientation even when tested with a first-order target (see Smith et al., 2001.))

When two contexts were matched based on the low-but not the high-level orientation content, the context with the higher-level orientations usually caused more bias when in competition (Img vs. Mixed in Fig. 9). Moreover, this effect was mediated by the task, indicating that the contexts were correctly balanced in terms of low-level statistics (see the “Detection task” in Fig. 9). Therefore, the spatial arrangement of the oriented content in the image affect the TAE. This result is consistent with modulation of TAE by perceptual grouping (He et al., 2012; Suzuki et al., 2005). Interestingly, the task dependence of the effect, and the lack of an effect in the absence of competition (compare “Img” to “Phs” in Fig. 9) are both consistent with a selection process between the different orientations, with preference in favor of the more task-relevant content (Ismail, Solomon, Hansard, & Mareschal, 2019; Pinchuk-Yacobi, Harris, et al., 2016). It is also interesting to note that visual attention, known to increase TAE magnitudes, seems to lead to a relatively modest and inconsistent effect (∼25%) (Spivey & Spirn, 2000). The large effect magnitudes observed here when the different contexts are overlapping, suggests that selection has more of an effect in the presence of strong competition.

### Contextual influence as a sum of separately bound contexts

Taken together, the results in this work showed an additive (linear) influence only when the contexts were separated. Here, we attempted to answer the question of why some contextual combinations are additive whereas others are not. Above, we identified two patterns of non-linearity: sub-additivity and selection. We argue that the findings of sub-additivity can be explained by compressive transformation of visual information during early stimulus processing, which binds visual stimulation using a narrow window of space and time into a higher-order structure. This idea is supported by accounts of contextual influence in terms of mechanisms exploiting statistical regularities in the visual environment (Coen-Cagli et al., 2015; Schwartz et al., 2007; Summerfield & De Lange, 2014). The contextual influence is then determined by selection between the competing processed contexts (e.g., between high vs. low alternatives). Moreover, the influence (bias in degrees) is linear, given a selected, processed context. To summarize, sufficiently separated contexts are bound separately, and will accumulate additively in terms of measured bias, weighted by selection:

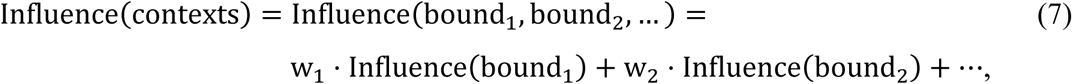

where w_i_ are the selection weights (possibly binary). A simple interpretation of these findings is that contextual influence and visual perception, both non-linear transformations, are at least partially shared during visual processing. This can be useful for the system in two ways. First, resolving an orientation from a stimulus is a computationally challenging task, so it seems reasonable to “recycle” the previously resolved orientations. Second, the information about the properties of an object becomes saturated when the object is fully resolved. For example, with a large enough grating, the orientation of the grating can be estimated to a sufficiently high accuracy, so an increase in grating size, contrast, or viewing duration will not provide much new information. However, a different object can be considered as a separate source of information, so its influence is not saturated. Overall, it seems possible that an efficient implementation of orientation context dependence relies on previously resolved orientations.

We note that here we investigated somewhat extreme cases of contextual “separation”: a large temporal offset (600 ISI) vs. narrow or no spatial offset (adjacent/overlapping). It therefore remains unclear whether shorter ISIs, or larger spatial separations, will measure additivity or not.

### Bayesian cue combination

A possible realization of the above idea was investigated using a Bayesian framework with independent Gaussian cues that are used to infer an internal reference (Fig. 3). The inferred reference is partially, but not fully, local, as evident in the measurement of non-retinotopic TAEs (Fig. 4). Importantly, using model fitting, we found that the predictions of the cue-combination model were indistinguishable from additivity for the relevant parameter range, accounting for the observation of additivity with temporally separated contexts (Fig. 4).

Interestingly, when the cues cannot be assumed to be independent, the weight of the correlated cues is reduced (Oruç et al., 2003). This suggests an explanation of the observed sub-additivity of adjacent, and hence, correlated, contexts (Fig. 6). A similar argument applies for the observed sub-additivity of increased adaptor contrast (Fig. 5 and Blake et al., 2006) or duration (Dekel & Sagi, 2015; Sekuler & Littlejohn, 1974) (see the discussion above). In the TAE experiments with temporally separated contexts, the correlation between cues was presumably negligible, which led to additivity. Note that estimating the relevant correlations using natural statistics is inaccurate, because reference cues probably change more slowly in space than is implied by image correlations (for example, two neighboring image regions can have very different image properties but the same frame of reference). Still, qualitatively, the results are in agreement with a faster rate of decay of TAE in foveal compared with peripheral vision (Fig. 4; Gao, Webster, & Jiang, 2019), consistent with temporal correlations in retinal images that may decay faster in fixation (Kayser, Einhäuser, & König, 2003; see also, Gekas et al., 2019; Snow, Coen-Cagli, & Schwartz, 2016). The derivation of quantitative estimations may require more detailed analysis.

Another prediction of the theory is that the influence of each cue depends on the ratio between the cue reliability and the unadapted vertical-reference reliability (see Eq. (6)) (the unadapted reference encompasses all the unknown unadapted reference cues). Based on this prediction, the TI and TAE are expected to be reduced when the unadapted reference is more reliable. For example, when a luminance frame of reference is present (see the appendix of Knapen et al., 2010).

It can also be observed that based on the model, bias is expected to be linear with the orientation difference between the context and the test (Eq. (6)). This prediction is not consistent with behavior: bias is known to reach a maximum for orientation differences of ∼20° (e.g., Schwartz et al., 2007). Still, this discrepancy should probably not be considered as a strong argument against the theory, because orientations that are sufficiently far from the reference are obviously not suited to serve as good reference cues (and hence deserve a lower weight). More extensive modeling is outside the scope of this work, but TAE may indeed be linear with small orientation differences (Campbell & Maffei, 1971; Magnussen & Kurtenbach, 1980).

Overall, the theory suggests how different reference cues are combined, which can account for the TAE and the TI. What counts as a reference cue may vary, including physical frames, predictions (Pinchuk-Yacobi, Dekel, et al., 2016), gravity (Wolfe & Held, 1982), and proprioception. In addition, the proposed theory probably does not account for all cases of orientation bias (for possible components of bias, see Clifford, Wenderoth, & Spehar, 2000). For example, the TAE experiments considered here use brief adaptor presentations (50 ms) and comparatively long inter-stimulus intervals (600 ms), a situation where contrast adaptation can be argued to be negligible. Possibly, other experimental situations could be explained better by contrast adaptation, which may be an additional, separate cause of the TAE.

### Orientation discrimination sensitivity (JND)

Across experiments, we measured the orientation discrimination sensitivity (JND, see Methods and Results). Overall, changes in JND by context were small (10-20%) and in most cases not statistically significant. This is perhaps not too surprising, given that the literature does not describe a universal relationship between bias and JND. For example, learning of one may affect the other, but the time course may differ (Chen & Fang, 2011). Still, under certain experimental manipulations, bias and JND may be positively correlated (Regan & Beverley, 1985; Solomon & Morgan, 2006). These effects are observed as increased JND in the presence of context as compared with its absence. Such a comparison is not available from our TI experiment, where a context was present in all experimental manipulations. These contextual variations were found to have no effect on JND. Regarding TAE, a no-context condition is available, indeed showing a smaller, though not statistically significant, JND as compared with the measurements in the presence of context (“empty” Gabor adaptors in Fig. 4E, and zero contrast image adaptors in Fig. 5B).

The cue combination theory (see Modeling and Figs. 3 and 4) does not have any predictions concerning JND. Note that cue reliability (Eq. 3) refers to the validity of using the adaptor/surround orientation as a cue for the true vertical, which is only partially influenced by its inter-trial stability, while JND depends on the inter-trial stability of the estimated vertical reference (Morgan, Watamaniuk, & McKee, 2000), and on orientation sensitivity of the sensory system. Therefore, the theory is agnostic about whether the inter-trial stability of the cues is better or worse than that of the unadapted reference. Consequently, in the presence of the adaptor/surround, the inter-trial stability of the reference can either improve or deteriorate. For example, our natural-image adaptors have much worse inter-trial stability than Gabor adaptors, consistent with the larger JNDs with image but not Gabor adaptors (Fig. 5B) (see also Erlikhman, Singh, Ghose, & Liu, 2019).

## Acknowledgments

This research was supported by the Basic Research Foundation, administered by the Israel Academy of Science. We thank Drs. Misha Katkov and Noga Pinchuk-Yacobi for their suggestions and comments.

## Data and code availability

The data that support the findings of this study, the images used in the behavioral experiments, and a workable example of the code used in the web experiments are all available from the corresponding author upon reasonable request.

